# Mildew Locus O facilitates colonization by arbuscular mycorrhiza in angiosperms

**DOI:** 10.1101/851840

**Authors:** Catherine N. Jacott, Myriam Charpentier, Jeremy D. Murray, Christopher J. Ridout

## Abstract

- Loss of barley *Mildew Resistance Locus O* (*MLO*) is known to confer durable and robust resistance to powdery mildew (*Blumeria graminis*), a biotrophic fungal leaf pathogen. Based on the increased expression of *MLO* in mycorrhizal roots and its presence in a clade of the MLO family that is specific to mycorrhizal-host species, we investigated the potential role of MLO in arbuscular mycorrhizal interactions.
- Using mutants from barley, wheat, and *Medicago truncatula*, we demonstrate a role for MLO in colonization by the arbuscular mycorrhizal fungi *Rhizophagus irregularis*.
- Early mycorrhizal colonization was reduced in *mlo* mutants of barley, wheat and *Medicago truncatula*, and this was accompanied by a pronounced decrease in the expression of many of the key genes required for intracellular accommodation of arbuscular mycorrhizal fungi.
- These findings suggest that the primary role of MLO in angiosperms is in the establishment of symbiotic associations with beneficial fungi, which has been appropriated by powdery mildew.

## Introduction

Plants have coevolved with microbes, which is evident in the elaborate strategies that plants use to both promote beneficial symbioses and to restrict pathogenesis. For example, to establish beneficial endosymbioses like arbuscular mycorrhization, plants have dedicated host signaling pathways involving hundreds of genes (Bonfante & Genre, 2010; Oldroyd, 2013; Bravo *et al.*, 2016). Similarly, to keep apace with pathogens, components of the plant immune system are rapidly evolving and expanding (Chisholm *et al.*, 2006; Jones & Dangl, 2006). Pathogens employ strategies to take advantage of host genes—so-called ‘susceptibility factors’—to cause disease (Panstruga, 2003; O’Connell & Panstruga, 2006). One example of a susceptibility factor is the barley *Mildew resistance locus O* (*MLO)* gene—referred to here as *MLO1* to distinguish it from other family members—which is required for successful colonization by powdery mildew fungi, *Erysiphales* (Jørgensen, 1992). However, the extent to which host susceptibility factors are required for beneficial symbioses is unknown.

MLO encodes a plasma membrane protein that possesses seven transmembrane helices and a C-terminal calmodulin-binding domain (Devoto *et al.*, 1999; Kim *et al.*, 2002). Most research on MLO1 has been on its role in susceptibility to powdery mildew. Knock-out mutants of *MLO1* in barley have been successfully deployed to provide broad-spectrum, durable resistance to powdery mildew since 1942 (Freisleben & Lein, 1942; Jørgensen, 1992). Despite considerable progress on the characterization of the powdery mildew resistance phenotype and its underlying molecular mechanism (Kim *et al.*, 2002; Piffanelli *et al.*, 2002; Consonni *et al.*, 2010), MLO1’s biological role remains unclear. Notably, orthologs of MLO1 are only present in those plant species that can host arbuscular mycorrhizal fungi (Bravo *et al.*, 2016). The study of other MLO family members has revealed their involvement in diverse plant processes, including pollen tube reception (AtMLO7/NORTIA) and root thigmomorphogenesis (Kessler *et al.*, 2010; Bidzinski *et al.*, 2014). The ancestral function of MLO proteins is unknown, but their presence in green algae suggests that the so-far characterized roles are derivative (Takamatsu, 2004; Jiao *et al.*, 2011). In *Arabidopsis thaliana*, loss of MLO family members belonging to a separate clade from barley MLO1 confer resistance to powdery mildew in a redundant fashion (Kusch *et al.*, 2016; Kuhn *et al.*, 2017). Interestingly, one of these can complement the *Atmlo7* pollen tube reception phenotype, suggesting a conserved biochemical function of MLO proteins (Jones *et al.*, 2017).

Most land plants can form an endosymbiosis with members of the *Glomeromycota*, reflecting the ancient origins of this interaction (Remy *et al.*, 1994; Parniske, 2008). This mutualistic relationship supplies water and nutrients—particularly phosphate, but also nitrogen and zinc— to the host plant (Cooper & Tinker, 1978; Lambert *et al.*, 1979; Govindarajulu *et al.*, 2005; Watts-Williams & Cavagnaro, 2012). Several key components of the signaling pathway required for establishing mycorrhizal symbioses (the common SYM pathway) are conserved in green algae, suggesting that land plant ancestors were preadapted for symbiosis (Delaux *et al.*, 2013; Delaux *et al.*, 2015). Furthermore, MLO1 was previously suggested to have a role in arbuscular mycorrhization (Ruiz-Lozano *et al.*, 1999). However, the analysis was limited to a single *mlo* allele, which was evaluated at a single time point. Since then, an additional study found no evidence supporting a role for an MLO1 homolog in arbuscular mycorrhization of pea (Humphry *et al.*, 2011).

Using mutants from barley, wheat, and *M. truncatula* and transcript profiling of *Hvmlo1* mycorrhizal roots, we reveal an unequivocal role for MLO in the early stages of arbuscular mycorrhizal colonization in angiosperms. The findings suggest that the primary role of MLO is in the establishment of fungal symbioses, which has been exploited by powdery mildew.

## Materials and methods

### Phylogenetic analyses

Sequences were aligned using MUSCLE (Edgar, 2004), and a maximum likelihood tree was constructed using MEGA X following a Jones-Taylor-Thornton (JTT) model with 100 bootstrap replications (Kumar *et al.*, 2018). Protein sequences and references are in Methods S1.

### Barley, wheat and M. truncatula mutants

Details of the barley and wheat *mlo* mutants used are shown in Methods S2. To identify homozygous *M. truncatula mlo8 Tnt1-*insertion mutants, DNA was extracted (DNeasy 96 Plant Kit, Qiagen). Genotyping of segregating seedling populations was performed by PCR using primers listed in Methods S3. To detect the *MtMLO8* cDNA, RNA was extracted from mycorrhizal *M. truncatula* roots (Plant RNeasy Kit, Qiagen). DNases were removed (TURBO DNA*-free*, Ambion), and 150 ng of RNA was retrotranscribed (SuperScript IV reverse transcriptase, Invitrogen). cDNA was amplified by RT-PCR (*Phusion* High Fidelity Polymerase kit, New England Biolabs) using primers listed in Methods S3.

### Plant growth conditions and mycorrhization and powdery mildew assays

For mycorrhization experiments, barley, wheat, and *M. truncatula* were grown in pots containing 80-90% Terragreen/sand and 10-20% mycorrhizal inoculum (homogenized soil substrate containing *Allium schoenoprasum* roots exposed to *Rhizophagus irregularis* for 8 weeks). Full descriptions of seed sterilization, soils substrates, and growth conditions for mycorrhization and powdery mildew assays are provided in Methods S4.

### Agrobacterium rhizogenes-mediated gene transfer

*A. rhizogenes*-mediated gene transfer (Boisson-Dernier *et al.*, 2001), was performed using strain AR1193. The promoter-GUS construct *pMtMLO8:GUS* was generated using Gateway® technology (Primrose & Twyman, 2013): entry vector pDONR207; destination vector 243_pKGW-GGRR. The 1983 bp promoter region of *MtMLO8* (Medtr3g115940) was amplified using primers with Gateway-compatible end sequences (Methods S3). The complementation construct *pMtMLO8:MtMLO8* was generated using the Golden Gate system (Engler *et al.*, 2009) using the same *MtMLO8* promoter region fused to the 1656 bp coding region. Domesticated DNA parts were synthesized by GeneArt® (Life Technologies). pL2V-50507 was used for the L2 backbone, with *pLjUBIQUTIN:DsRed:t53S* placed in position 1, and *pMtMLO8:MLO8:t35S* in position 2.

### Fungal staining and quantification

Staining (ink, GUS, and WGA-Alexa488) and visualization methods are summarized in Methods S5. Samples were blinded and mycorrhizal infection structures quantified using the gridline intersect method (Giovannetti and Mosse, 1980). Powdery mildew fungal structures were quantified by calculating the ratio of germinated spores: colony formation.

### Gene expression analyses

RNAs were extracted from root tissue (Plant RNeasy Kit, Qiagen) and was treated with DNase (RNase-Free DNase Set, Qiagen). Methods for RT-qPCR and transcriptome analyses can be found in Methods S6.

## Results

To assess a potential function of MLO1 in mycorrhization, we compared expression levels of barley *MLO1* (*HvMLO1*) in mycorrhizal and non-mycorrhizal roots (**Fig. 1a**). At 22 days post-inoculation (dpi) with *R. irregularis*, RT-qPCR analysis indicated that *HvMLO1* was induced in mycorrhizal roots. We then evaluated mycorrhizal phenotypes of barley *Hvmlo1* mutants by quantification of mycorrhizal fungal infection structures over a time-course. Arbuscules and vesicles were quantified at 22, 29, and 36 dpi with *R. irregularis* (20% inoculum) in barley cv. Ingrid wild type, *Hvmlo1-1*, and *Hvmlo1-5* roots (**Fig. 1b**). We found a reduction in arbuscule and vesicle occurrence in *Hvmlo1-1* and *Hvmlo1-5* compared to wild type at 22 dpi. This difference was maintained in *Hvmlo1-5* at 29 dpi, whereas *Hvmlo1-1* showed a similar phenotype to wild type. By 36 dpi, no significant difference in mycorrhization between the lines was detectable. Wild type roots appeared to have established full mycorrhization between 22 and 29 dpi, whereas the *Hvmlo1* mutants established full colonization between 29 and 36 dpi. Overall, reduced mycorrhization was more evident at early time points, suggesting that mycorrhizal development in *Hvmlo1* is delayed, but later recovers to wild type levels.

**Fig. 1.**
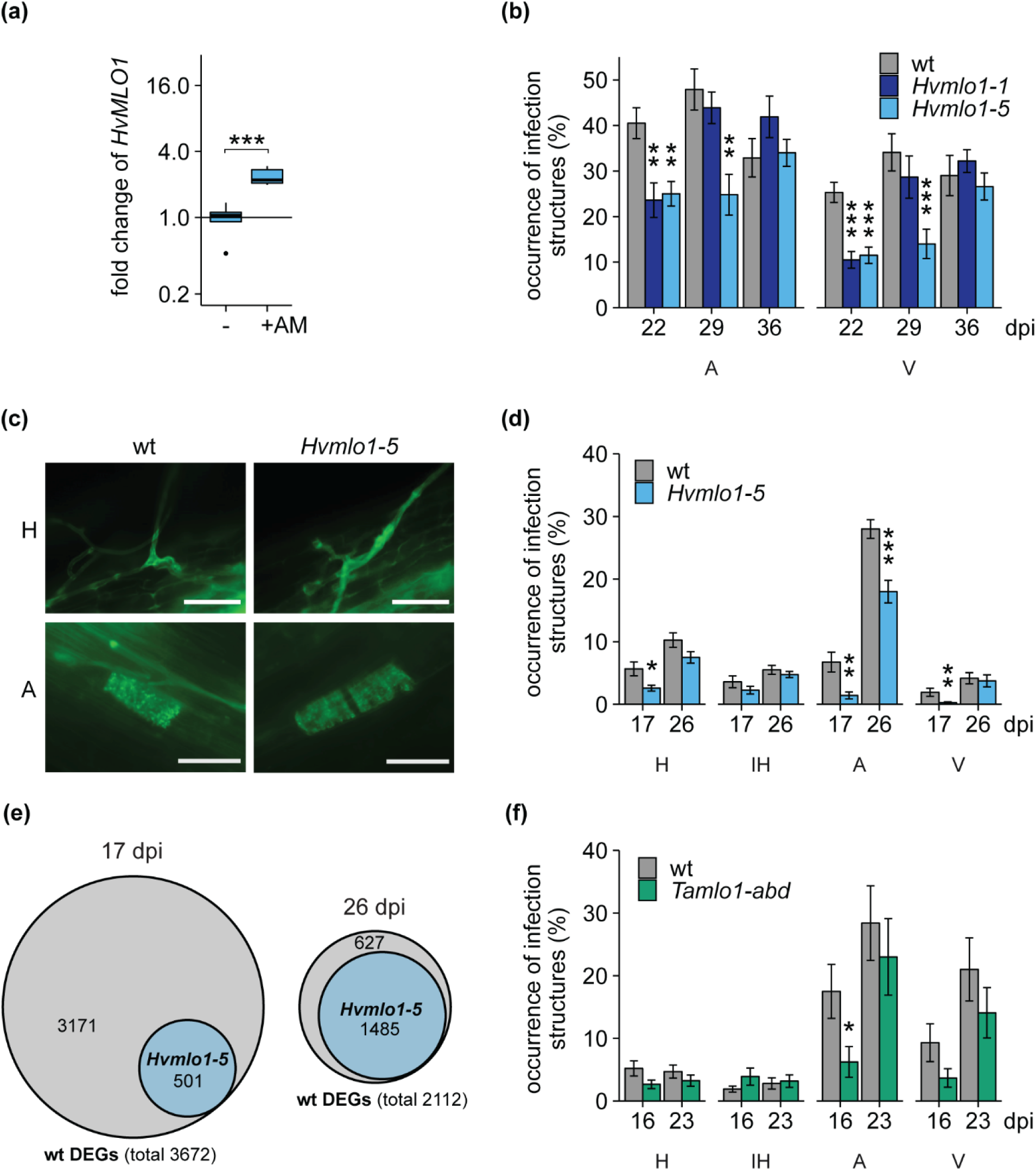
Mycorrhizal colonization in barley and wheat *mlo1* mutants. **(a)** Relative expression of *HvMLO1* in wild type barley cv. Ingrid roots without (-) and with *R. irregularis* (+AM) at 22 dpi. Relative expression levels were measured by RT-qPCR and normalized to *HvEF1alpha* (barley). Statistical comparisons were made relative to non-mycorrhized root samples. Bars represent means of 8 biological replicates ± SEM (Student’s *t*-test: ***P <0.001). **(b, d, f)** Quantification of arbuscular mycorrhizal infection structures in barley cv. Ingrid and wheat cv. KN1999 wild type and *mlo1* roots after inoculation with *R. irregularis*: hyphopodia (H), intraradical hyphae (IH), arbuscules (A), and vesicles (V). The binomial occurrences of mycorrhizal structures are shown as a percentage of the total number of root sections assessed. Statistical comparisons have been made to the wild type. Values are the mean of 12 biological replicates ±SEM (error bars) (General Linear Model with a logit link function; ANOVA; *, P < 0.05; **, P < 0.01; ***, P < 0.001). **(c)** Appearance of hyphopodia (H) and arbuscules (A) in wild type and *Hvmlo1-5* roots at 17 dpi with *R. irregularis*. Mycorrhizal fungal structures were stained using Alexa Fluor 488 wheat germ agglutinin. Photos indicate representative examples from 40 observations. Scale bars = 50 μm. **(e)** Venn diagrams showing the number of differentially regulated genes (DEGs) in mycorrhizal roots relative to uninoculated roots in wild type and the proportion of genes similarly responding in *Hvmlo1-5* using a cut-off of fold change > 2; FDR-corrected P-value < 0.05.

To further assess the mycorrhizal phenotype of *Hvmlo1* during the early stages of mycorrhization, we performed a detailed analysis of symbiotic structures (hyphopodia, intraradical hyphae, arbuscules and vesicles) at early time points using less *R. irregularis* inoculum (10% inoculum) (**Fig. 1c,d**). At 17 dpi, *Hvmlo1-5* showed a statistically significant reduction in the occurrence of hyphopodia, arbuscules, and vesicles compared to wild type. Similar to the previous experiment, by 26 dpi, *Hvmlo1-5* showed a statistically significant reduction in only arbuscule occurrence compared to wild type. A similar phenotype was observed between wild type and *Hvmlo1-5* in a different genetic background, barley cv. Pallas, in two separate experiments (**Fig. S1**). We observed no differences in the physical structure of hyphopodia or arbuscules in *Hvmlo1-5* mutants (**Fig. 1c**).

To gain insight into the nature of the delayed infection in *Hvmlo1-5*, we carried out comparative transcriptomic analyses at 17 dpi and 26 dpi with *R. irregularis* (**Fig. 1e**). At 17 dpi, there were considerably fewer differentially regulated genes (DEGs) in *Hvmlo1-5* than wild type, with 86% (3171/3672) of all wild type DEGs not responding in *Hvmlo1-5.* Whereas by 26 dpi, despite the recovery of observable colonization in the mutant, there was still a substantial (∼30%) reduction of the number of DEGs in *Hvmlo1-5* relative to wild type, suggesting persistent perturbation of the mycorrhizal interaction.

To further investigate the nature of the differences in *Hvmlo1-5*, we identified the sequences of potential barley orthologues of known mycorrhizal genes previously identified in other species and examined their expression in our data. Amongst the genes induced more than 2-fold in wild type—but not in *Hvmlo1-5*—at 17 dpi, were genes required for early infection and components of the common signalling pathway required for nodulation and mycorrhization, *VAPYRIN, NSP1, NSP2, DMI2, IPD3* and *NOPE1* homologues (Catoira *et al.*, 2000; Ané *et al.*, 2002; Lévy *et al.*, 2004; Messinese *et al.*, 2007; Pumplin *et al.*, 2010; Murray *et al.*, 2011; Nadal *et al.*, 2017) (**Table 1, Table S1-S3, Fig. S2**). This suggests that *MLO1* is required for timely or full-activation of early processes of mycorrhizal colonization. In addition, potential orthologues of several genes involved in arbuscule development and function were affected in *Hvmlo1-5*, including *RAM1, RAD1, RAM2, EXO70I, STR, STR2 AMT2-3, AMT3-4* and *PT4* homologues (Harrison *et al.*, 2002; Zhang *et al.*, 2010; Gobbato *et al.*, 2012; Wang *et al.*, 2012; Breuillin-Sessoms *et al.*, 2015; Xue *et al.*, 2015; Zhang *et al.*, 2015). Generally, these genes were induced to a lesser extent than wild type at 17 dpi, and partly or fully recovered by 26 dpi, which mirrors the initial delay and later recovery of the colonization phenotype. Together, these phenotypic and transcriptomic results indicate a role for MLO1 in the early stage of mycorrhization.

**Table 1:**
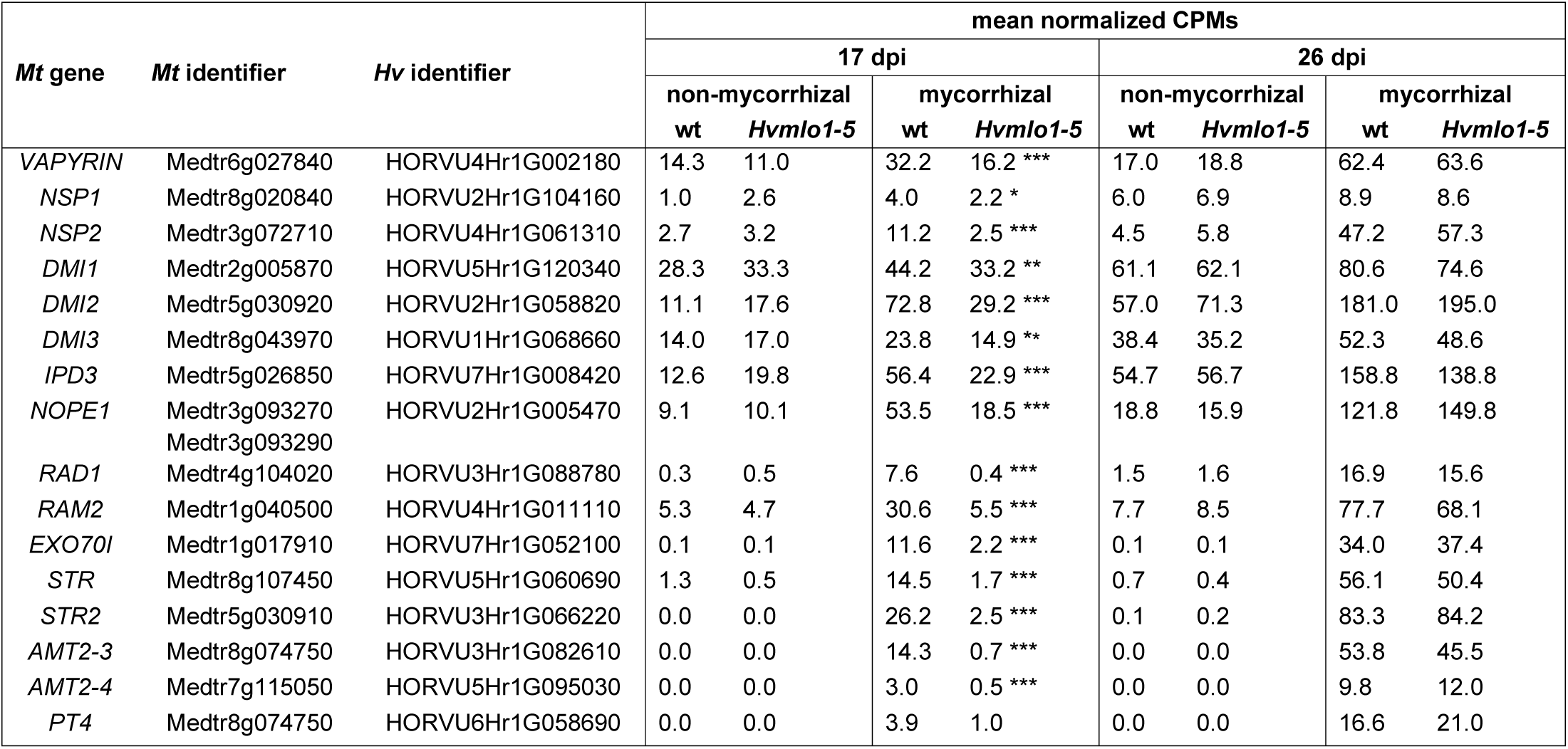
Counts per million (CPM) values of potential orthologues of previously described mycorrhizal genes in wild type and *Hvmlo1-5* non-mycorrhizal and mycorrhizal roots: significant fold changes between wt and *Hvmlo1-5* at corresponding time points and treatments are indicated (FDR-corrected P-value; *, P < 0.05; **, P < 0.01; ***, P < 0.001).

To investigate the conservation of function for this MLO, we used phylogenetic analysis to identify MLO proteins which may be functionally analogous to barley HvMLO1 (**Fig. 2**). The analysis included mycorrhizal host species: basal angiosperm *Amborella trichopoda* (Atr); dicots *Glycine max* (Gm), *M. truncatula* (Mt), *Pisum sativum* (Ps) and *Solanum lycopersicum* (Sl); and monocots *H. vulgare* (Hv), *Oryza sativa* (Os) and *Triticum aestivum* (Ta), as well as non-mycorrhizal host species: moss *Physcomitrella patens* (Pp); and dicots *A. thaliana* (At), *Beta vulgaris* (Bv), *Dianthus caryophyllus* (Dc) and *Lupinus angustifolius* (La). As previously shown (Kusch *et al.*, 2016), the phylogenetic analysis supported the view that embryophyte MLO proteins diverge into seven clades. HvMLO1 groups in clade IV, which is comprised only of species that are mycorrhizal-hosts, while powdery mildew susceptibility factors identified in monocots and dicots are found in clades IV and V, respectively.

**Fig. 2.**
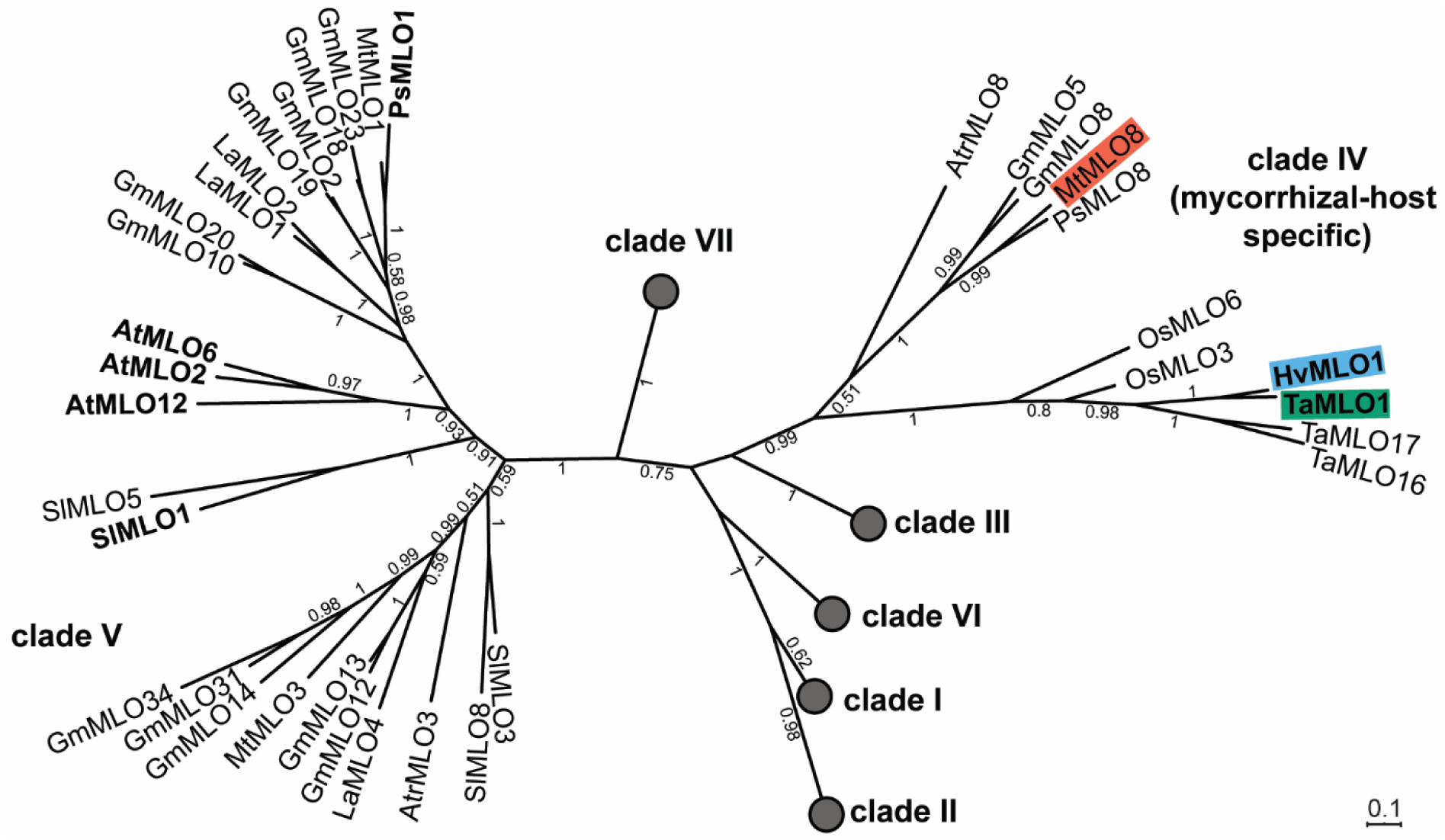
MLO phylogeny. A phylogenetic tree showing the relationship between MLO proteins from *A. trichopoda, A. thaliana, B. vulgaris, D. caryophyllus, G. max, H. vulgare, L. angustifolius, M. truncatula, O. sativa, P. sativum, P. patens, S. lycopersicum and T. aestivum.* The *T. aestivum* MLO protein dataset often included near-identical homoeoalleles (A, B, and D copies); therefore, for simplicity, only one homoeoallele was included in the phylogenetic analysis (A homoeoallele where possible). The tree was generated using MEGA X from an amino acid alignment. The phylogenetic tree was calculated via the maximum likelihood method, using JTT modeling with 100 bootstrap replications. Bootstrap values > 0.5 are shown. The scale bar indicates the evolutionary distance based on the amino acid substitution rate. Circles indicate collapsed clades; known powdery mildew susceptibility factors are labeled in bold, and candidate MLO proteins for a role in mycorrhization are highlighted.

The wheat functional orthologue of HvMLO1, TaMLO1, is a powdery mildew host susceptibility factor (Wang *et al.*, 2014). *Tamlo1-abd* contains TALEN-induced mutations in all three homoeoalleles (A, B, and D) (Wang *et al.*, 2014). To assess the mycorrhizal phenotype of *Tamlo1-abd* during the early stages of mycorrhization, we performed a detailed analysis of mycorrhizal structures (hyphopodia, intraradical hyphae, arbuscules, and vesicles) at early time points similar to those used for barley (**Fig. 1f**). At 16 dpi with *R. irregularis, Tamlo1-abd* showed a significant reduction in arbuscule occurrence compared to wild type. However, consistent with our findings in barley, by 23 dpi *Tamlo1-abd* colonization levels were similar to wild type. Wheat *mlo1* mutants exhibited the delayed mycorrhizal phenotype in two separate experiments, suggesting a conserved role for MLO1 in mycorrhization in cereals.

To determine whether MLO’s role in mycorrhization extends to dicots, we investigated *M. truncatula* MtMLO8, the apparent orthologue of HvMLO1, which was previously found to be conserved exclusively in mycorrhizal host plants (Bravo *et al.*, 2016). We first tested its expression in mycorrhizal roots, and we found that like *HvMLO1, MtMLO8* was induced relative to non-mycorrhizal roots (**Fig. 3a, Fig. S3a**), consistent with data from several studies available in the public database (Benedito *et al.*, 2008; Breakspear *et al.*, 2014; Luginbuehl *et al.*, 2017). To establish in which cell types *MtMLO8* is expressed, we studied its expression in *M. truncatula* roots using the *MtMLO8* promoter to drive *GUS* expression using the *Agrobacterium rhizogenes* mediated hairy root transformation system. *MtMLO8* promoter activity was strongly associated with cortical cells containing arbuscules (**Fig. 3a, S3b**).

**Fig. 3.**
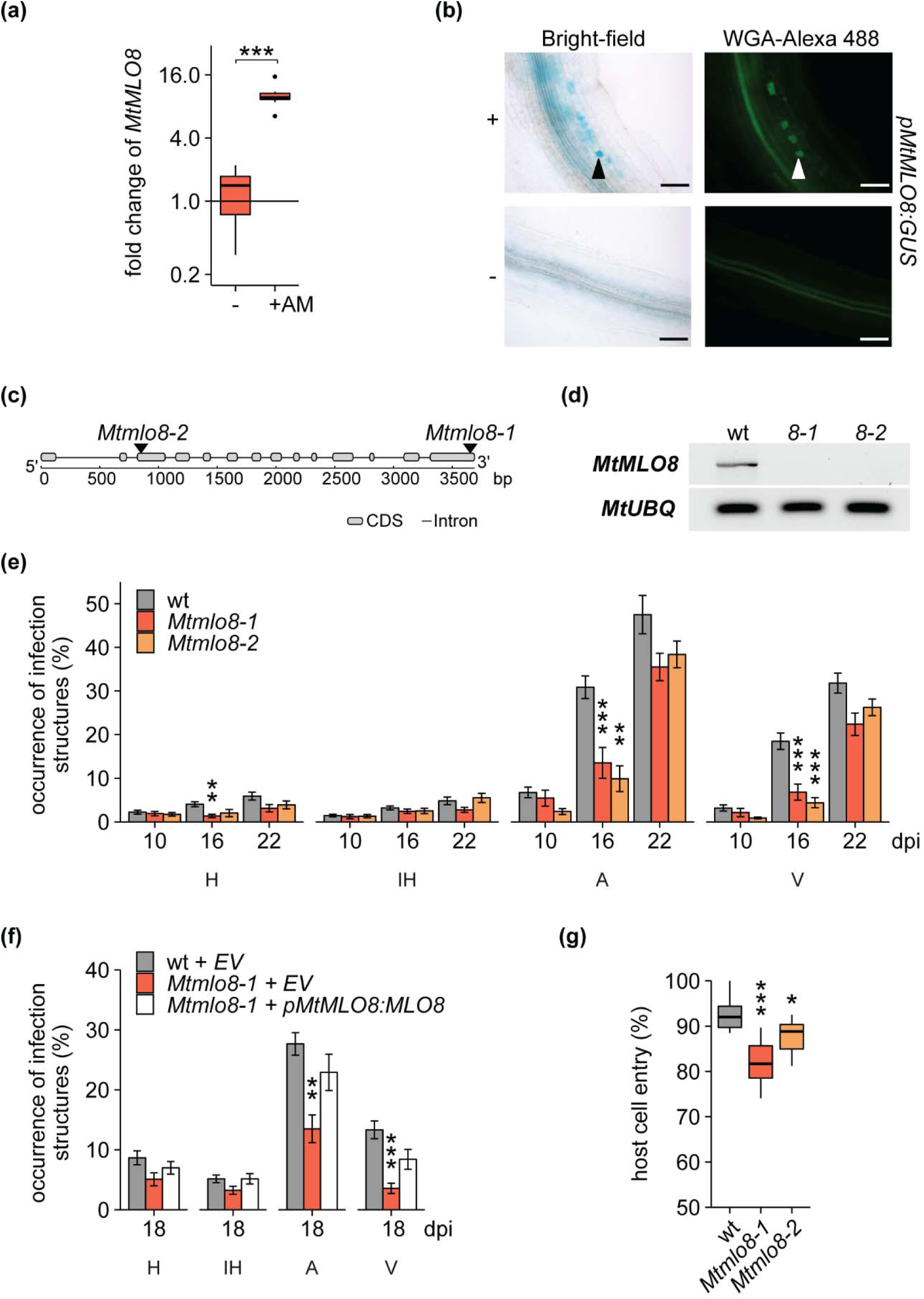
MtMLO8 is involved in mycorrhizal colonization of *M. truncatula.* **(a)** Relative expression of *MtMLO8* in wild type *M. truncatula* cv. R108 without (-) and with *R. irregularis* (+AM) at 22 dpi. Relative expression levels were measured by RT-qPCR and normalized to the geometric mean of *MtUBIQUITIN* and *MtPTB*. Statistical comparisons were made relative to non-mycorrhized root samples. Bars represent means of 8 biological replicates ± SEM (error bars) (Student’s *t*-test: ***P <0.001). **(b)** Activity of the *MtMLO8* promoter in mycorrhized and non-mycorrhized wild type roots at 21 dpi with *R. irregularis*, assessed using a promoter-GUS fusion. Bright-field and corresponding green fluorescence images of *M. truncatula* hairy roots expressing the β-*glucuronidase* (*GUS*) gene under the control of the *MtMLO8* promoter. Mycorrhizal fungal structures were visualized using Alexa Fluor 488 wheat germ agglutinin. Solid arrowheads indicate cells containing arbuscules and empty arrowheads indicate hyphopodia. Scale bars = 100 μm. **(c)** Gene structure of *M. truncatula MtMLO8*. Arrows indicate the *Tobacco retrotransposon 1* (*Tnt1*) insertion sites in the *Mtmlo8* mutants. **(d)** RT-PCR was used to detect the accumulation of the *MtMLO8* transcript in wild type, *Mtmlo8-1*, and *Mtmlo8-2* mutant roots. *MtUBIQUITIN* was used as a constitutive control. **(e)** Quantification of mycorrhizal structures in *M. truncatula* cv. R108 wild type (wt), *Mtmlo8-1* and *Mtmlo8-2* roots. **(f)** Complementation of the *Mtmlo8-1* mutants and quantification of mycorrhizal structures in hairy roots. *Mtmlo8-1* was transformed with *pMtMLO8:MtMLO8* or empty vector control (EV), and wild type were transformed with EV control. The binomial occurrences of mycorrhizal structures, hyphopodia (H), intraradical hyphae (IH), arbuscules (A) and vesicles (V), are shown as a percentage of the total number of root systems assessed. **(g)** Quantification of powdery mildew infection on wild type, *Mtmlo8-1* and *Mtmlo8-2* leaves at 72 hpi with *Erysiphe pisi*. The binomial occurrences of successful host cell entry are shown as a percentage of the total number of spores observed. For figures **e, f, g** statistical comparisons have been made to the wild type. Bars represent the mean of 15 (d), 10 (e) and 10 (f) biological replicates ± SEM (error bars) (General Linear Model with a logit link function; ANOVA, post hoc pairwise; *, P < 0.05; **, P < 0.01; ***, P < 0.001).

To investigate whether MtMLO8 functions during mycorrhization, seeds from two independent *M. truncatula* cv. R108 lines carrying *Tnt1* insertions in the coding sequence of these genes were obtained, and homozygous mutants were identified (**Fig. 3c, d**) (Tadege *et al.*, 2008). We performed a detailed analysis of mycorrhizal structures (hyphopodia, intraradical hyphae, arbuscules, and vesicles) at multiple time points in wild type, *Mtmlo8-1* and *Mtmlo8-2*. Compared to wild type, there was a significant reduction in arbuscules and vesicles at 16 dpi in both alleles of *Mtmlo8* (**Fig. 3e**), consistent with our results using the orthologous mutants in barley and wheat. By 22 dpi—and at later time points (**Fig. S4a**)—there was no difference between the lines. We observed no impairment in the physical structure of mycorrhizal structures, hyphopodia and arbuscules in *Mtmlo8* mutants (**Fig. S4b**). To validate that mycorrhizal phenotype observed was due to mutations in *MtMLO8*, and not a consequence of additional *Tnt1* insertions or other mutations in the mutant backgrounds, we expressed *MtMLO8* from its native promoter in *Mtmlo8-1* mutant roots by transformation with *A. rhizogenes.* When arbuscule occurrence in roots of wild type plants transformed with the empty vector reached approximately 30%—corresponding to when the *Mtmlo8* phenotype could be observed—mycorrhizal structures (hyphopodia, intraradical hyphae, arbuscules, and vesicles) were quantified in the transgenic roots. The expression of *MtMLO8* under the control of its native promoter successfully complemented the *Mtmlo8-1* mutant phenotype (**Fig. 3f**). In summary, these results suggest that mutations in *MtMLO8* affect the early stages of mycorrhization and delay arbuscule development.

To assess whether, like HvMLO1 and TaMLO1, MtMLO8 might also have a role in powdery mildew susceptibility, we inoculated wild type and *Mtmlo8* mutant leaves with an isolate of powdery mildew that can infect *M. truncatula, Erysiphe pisi.* We observed a small but statistically significant reduction in the percentage of successful host cell entry (ratio of germinated spores: colony formation) in *Mtmlo8* compared to wild type (**Fig. 3g**). This result indicates that *MtMLO8* is required not only for early mycorrhizal infection but also for powdery mildew colonization.

## Discussion

Since MLO’s function in facilitating powdery mildew infection is disadvantageous to the host, it follows that it must also fulfill some other positive role that explains its conservation. The phylogenetic analyses, mutant phenotypes, and gene expression studies presented here point to a symbiotic role for members of the MLO clade IV and suggest that—at least in part— selection to maintain this gene is related to its function in mycorrhizal interactions. A common thread between the established roles of MLO in fungal interactions, pollen-tube reception, and thigmotropism is touch-sensing. Indeed, MLO proteins have been reported to accumulate at the site of powdery mildew penetration (Bhat *et al.*, 2005), as well as at the contact point between the pollen tube and synergid cell (Kessler et al., 2010). These responses could result from a generic response to mechanical stimuli. Notably, a role for mechanosensing has been proposed for the receptor kinase Feronia (Hamant & Haswell, 2017), which was shown to be required for proper re-localization of MLO7/NORTIA to the site of pollen tube contact in the synergid cell (Kessler *et al.*, 2010; Jones *et al.*, 2017). However, other cues may activate MLO, for example, both *Blumeria graminis* and *R. irregularis* produce effectors that could target MLO (Kloppholz *et al.*, 2011; Requena *et al.*, 2018; Pennington *et al.*, 2019; Saur *et al.*, 2019). Regardless of how it is activated, it will be of interest to study the localization of MLO1 or its orthologues during hyphopodium formation.

The mycorrhizal phenotype of the *mlo* mutants examined was not severe, but may be expected to have measurable consequences on fitness in a natural setting with competition for limited resources. Furthermore, global gene expression was still affected at the later time point, so a detailed study of growth and nutrient uptake characteristics of the mutant may show differences. In addition, the increased expression of other MLOs during mycorrhization (**Fig. S3a**) in *M. truncatula* suggests that—as for other *mlo* phenotypes such as powdery mildew resistance (Kuhn *et al.*, 2017) and synergid cell reception (Jones & Kessler, 2017)—MLOs act redundantly, with different MLOs potentially contributing unequally across different cell types and developmental stages. It is, therefore, possible that higher-order mutants will have increasingly stronger arbuscular mycorrhizal phenotypes. The powdery mildew phenotype of *Mtmlo8* was relatively minor. It is possible that multiple *MLO* genes contribute to powdery mildew susceptibility as is the case in *A. thaliana*, for example, *MtMLO1*—the apparent orthologue of *PsMLO*— which is involved in powdery mildew susceptibility in pea (Humphry *et al.*, 2011; Pavan *et al.*, 2011). Notably, the narrow time window for mycorrhizal phenotype definition and potential functional redundancy with gene family members could explain why MLO has not so far been identified in mutagenesis screens.

Despite having small yield penalties, mutants for MLO have been deployed in agriculture for decades. Understanding the role of MLO in symbiotic interactions with mycorrhizal fungi is therefore vital information for sustainable agriculture (Martin *et al.*, 2018). Our detailed phenotypic and transcriptomic analyses in both cereals and *M. truncatula* provide the basis for further elucidation of MLO function in mycorrhization and how plants balance this beneficial association with susceptibility to parasitic powdery mildews. As mycorrhizal and powdery mildew respectively infect root and shoot, it may be possible to generate genotypes that could fully support mycorrhiza while remaining non-hosts for powdery mildew.

## Acknowledgments

The wheat *Tamlo1-abd* was kindly supplied by Caixia Gao, State Key Laboratory of Plant Cell and Chromosome Engineering, Institute of Genetics and Developmental Biology, Chinese Academy of Sciences, Beijing China. *Erysiphe pisi* (Ep) CJ001 spores kindly supplied by Professor Diego Rubiales, Institute for Sustainable Agriculture, CSIC, Córdoba, Spain. The authors would like to thank Burkhard Steuernagel for aligning our RNA-sequencing data and advice with analyses; James Brown for advice on statistics; Alina-Andrada Igna for her preliminary experiments in *M. truncatula*; and Erin Zess for feedback during the editing of this manuscript. We want to thank past and present members of the Ridout, Murray, and Charpentier labs for help and advice. Jeremy Murray was supported by the National Key R&D Program of China (2016YFA0500500), the Biotechnology and Biological Sciences Research Council (grant no. BB/L010305/1 [David Phillips Fellowship]). Catherine Jacott was supported by the Biotechnology and Biological Sciences Research Council (BBSRC) Norwich Research Park Bioscience Doctoral Training Grants BB/J014524/1 and BB/M011216/1. Myriam Charpentier is supported by the BBSRC grant BB/P007112/1. Christopher J. Ridout is supported by BBSRC grant BB/P012574/1.

## Author contribution

CNJ performed experimental work and data analyses. MC, JDM, and CJR supervised and co-directed the research. CNJ, JDM, CJR and MC wrote the manuscript.

## References

Ané J-M, Lévy J, Thoquet P, Kulikova O, de Billy F, Penmetsa V, Kim D-J, Debellé F, Rosenberg C, Cook DR, et al. 2002. Genetic and Cytogenetic Mapping of *DMI1, DMI2,* and *DMI3* Genes of *Medicago truncatula* Involved in Nod Factor Transduction, Nodulation, and Mycorrhization. Molecular Plant-Microbe Interactions 15(11): 1108–1118.

Benedito VA, Torres-Jerez I, Murray JD, Andriankaja A, Allen S, Kakar K, Wandrey M, Verdier J, Zuber H, Ott T. 2008. A gene expression atlas of the model legume *Medicago truncatula*. The Plant Journal 55(3): 504–513.

Bhat RA, Miklis M, Schmelzer E, Schulze-Lefert P, Panstruga R. 2005. Recruitment and interaction dynamics of plant penetration resistance components in a plasma membrane microdomain. Proceedings of the National Academy of Sciences 102(8): 3135–3140.

Bidzinski P, Noir S, Shahi S, Reinstaedler A, Gratkowska DM, Panstruga R. 2014. Physiological characterization and genetic modifiers of aberrant root thigmomorphogenesis in mutants of *Arabidopsis thaliana MILDEW LOCUS O* genes. Plant, Cell & Environment 37(12): 2738–2753.

Boisson-Dernier A, Chabaud M, Garcia F, Bécard G, Rosenberg C, Barker DG. 2001. *Agrobacterium rhizogenes*-Transformed Roots of *Medicago truncatula* for the Study of Nitrogen-Fixing and Endomycorrhizal Symbiotic Associations. Molecular Plant-Microbe Interactions 14(6): 695–700.

Bonfante P, Genre A. 2010. Mechanisms underlying beneficial plant–fungus interactions in mycorrhizal symbiosis. Nature communications 1: 48.

Bravo A, York T, Pumplin N, Mueller LA, Harrison MJ. 2016. Genes conserved for arbuscular mycorrhizal symbiosis identified through phylogenomics. Nature Plants 2: 15208.

Breakspear A, Liu C, Roy S, Stacey N, Rogers C, Trick M, Morieri G, Mysore KS, Wen J, Oldroyd GE. 2014. The root hair “infectome” of *Medicago truncatula* uncovers changes in cell cycle genes and reveals a requirement for auxin signaling in rhizobial infection. The Plant Cell 26(12): 4680–4701.

Breuillin-Sessoms F, Floss DS, Gomez SK, Pumplin N, Ding Y, Levesque-Tremblay V, Noar RD, Daniels DA, Bravo A, Eaglesham JB, et al. 2015. Suppression of Arbuscule Degeneration in *Medicago truncatula phosphate transporter4* Mutants Is Dependent on the Ammonium Transporter 2 Family Protein AMT2;3. The Plant Cell 27(4): 1352–1366.

Catoira R, Galera C, de Billy F, Penmetsa RV, Journet E-P, Maillet F, Rosenberg C, Cook D, Gough C, Dénarié J. 2000. Four Genes of *Medicago truncatula* Controlling Components of a Nod Factor Transduction Pathway. The Plant Cell 12(9): 1647–1665.

Chisholm ST, Coaker G, Day B, Staskawicz BJ. 2006. Host-microbe interactions: shaping the evolution of the plant immune response. Cell 124(4): 803–814.

Consonni C, Bednarek P, Humphry M, Francocci F, Ferrari S, Harzen A, Ver Loren van Themaat E, Panstruga R. 2010. Tryptophan-derived metabolites are required for antifungal defense in the Arabidopsis *mlo2* mutant. Plant Physiology 152(3): 1544–1561.

Cooper KM, Tinker P. 1978. Translocation and transfer of nutrients in vesicular-arbuscular mycorrhizas: ii. Uptake and translocation of phosphorus, zinc and sulphur. New Phytologist 81(1): 43–52.

Delaux P-M, Radhakrishnan GV, Jayaraman D, Cheema J, Malbreil M, Volkening JD, Sekimoto H, Nishiyama T, Melkonian M, Pokorny L. 2015. Algal ancestor of land plants was preadapted for symbiosis. Proceedings of the National Academy of Sciences 112(43): 13390–13395.

Delaux P-M, Séjalon-Delmas N, Bécard G, Ané J-M. 2013. Evolution of the plant–microbe symbiotic ‘toolkit’. Trends in Plant Science 18(6): 298–304.

Devoto A, Piffanelli P, Nilsson I, Wallin E, Panstruga R, von Heijne G, Schulze-Lefert P. 1999. Topology, subcellular localization, and sequence diversity of the Mlo family in plants. Journal of Biological Chemistry 274(49): 34993–35004.

Edgar RC. 2004. MUSCLE: multiple sequence alignment with high accuracy and high throughput. Nucleic Acids Research 32(5): 1792–1797.

Engler C, Gruetzner R, Kandzia R, Marillonnet S. 2009. Golden Gate Shuffling: A One-Pot DNA Shuffling Method Based on Type IIs Restriction Enzymes. PloS one 4(5): e5553.

Freisleben R, Lein A. 1942. Über die Auffindung einer mehltauresistenten Mutante nach Röntgenbestrahlung einer anfälligen reinen Linie von Sommergerste. Naturwissenschaften 30(40): 608–608.

Gobbato E, Marsh JF, Vernié T, Wang E, Maillet F, Kim J, Miller JB, Sun J, Bano SA, Ratet P. 2012. A GRAS-type transcription factor with a specific function in mycorrhizal signaling. Current Biology 22(23): 2236–2241.

Govindarajulu M, Pfeffer PE, Jin H, Abubaker J, Douds DD, Allen JW, Bücking H, Lammers PJ, Shachar-Hill Y. 2005. Nitrogen transfer in the arbuscular mycorrhizal symbiosis. Nature 435(7043): 819–823.

Hamant O, Haswell ES. 2017. Life behind the wall: sensing mechanical cues in plants. BMC Biology 15(1): 59.

Harrison MJ, Dewbre GR, Liu J. 2002. A phosphate transporter from *Medicago truncatula* involved in the acquisition of phosphate released by arbuscular mycorrhizal fungi. The Plant Cell 14(10): 2413–2429.

Humphry M, Reinstädler A, Ivanov S, Bisseling T, Panstruga R. 2011. Durable broad-spectrum powdery mildew resistance in pea *er1* plants is conferred by natural loss-of-function mutations in *PsMLO1*. Molecular Plant Pathology 12(9): 866–878.

Jiao Y, Wickett NJ, Ayyampalayam S, Chanderbali AS, Landherr L, Ralph PE, Tomsho LP, Hu Y, Liang H, Soltis PS, et al. 2011. Ancestral polyploidy in seed plants and angiosperms. Nature 473(7345): 97–100.

Jones DS, Kessler SA. 2017. Cell type-dependent localization of MLO proteins. Plant Signaling & Behavior 12(11): 172–185.

Jones DS, Yuan J, Smith BE, Willoughby AC, Kumimoto EL, Kessler SA. 2017. MILDEW RESISTANCE LOCUS O Function in Pollen Tube Reception Is Linked to Its Oligomerization and Subcellular Distribution. Plant Physiology 175(1): 172–185.

Jones JD, Dangl JL. 2006. The plant immune system. Nature 444(7117): 323–329.

Jørgensen IH. 1992. Discovery, characterization and exploitation of Mlo powdery mildew resistance in barley. Euphytica 63(1-2): 141–152.

Kessler SA, Shimosato-Asano H, Keinath NF, Wuest SE, Ingram G, Panstruga R, Grossniklaus U. 2010. Conserved molecular components for pollen tube reception and fungal invasion. Science 330(6006): 968–971.

Kim MC, Panstruga R, Elliott C, Müller J, Devoto A, Yoon HW, Park HC, Cho MJ, Schulze-Lefert P. 2002. Calmodulin interacts with MLO protein to regulate defence against mildew in barley. Nature 416(6879): 447–451.

Kloppholz S, Kuhn H, Requena N. 2011. A Secreted Fungal Effector of *Glomus intraradices* Promotes Symbiotic Biotrophy. Current Biology 21(14): 1204–1209.

Kuhn H, Lorek J, Kwaaitaal M, Consonni C, Becker K, Micali C, Ver Loren van Themaat E, Bednarek P, Raaymakers TM, Appiano M. 2017. Key components of different plant defense pathways are dispensable for powdery mildew resistance of the Arabidopsis *mlo2 mlo6 mlo12* triple mutant. Frontiers in Plant Science 8: 1006.

Kumar S, Stecher G, Li M, Knyaz C, Tamura K. 2018. MEGA X: Molecular Evolutionary Genetics Analysis across Computing Platforms. Molecular Biology and Evolution 35(6): 1547–1549.

Kusch S, Pesch L, Panstruga R. 2016. Comprehensive phylogenetic analysis sheds light on the diversity and origin of the MLO family of integral membrane proteins. Genome Biology and Evolution 8(3): 878–895.

Lambert D, Baker DE, Cole H. 1979. The Role of Mycorrhizae in the Interactions of Phosphorus with Zinc, Copper, and Other Elements. Soil Science Society of America Journal 43(5): 976–980.

Luginbuehl LH, Menard GN, Kurup S, Van Erp H, Radhakrishnan GV, Breakspear A, Oldroyd GED, Eastmond PJ. 2017. Fatty acids in arbuscular mycorrhizal fungi are synthesized by the host plant. Science 356: 1175–1178.

Lévy J, Bres C, Geurts R, Chalhoub B, Kulikova O, Duc G, Journet E-P, Ané J-M, Lauber E, Bisseling T, et al. 2004. A Putative Ca2+ and Calmodulin-Dependent Protein Kinase Required for Bacterial and Fungal Symbioses. Science 303(5662): 1361–1364.

Martin FM, Harrison MJ, Lennon S, Lindahl B, Öpik M, Polle A, Requena N, Selosse MA. 2018. Cross-scale integration of mycorrhizal function. New Phytologist 220(4): 941–946.

Messinese E, Mun J-H, Yeun LH, Jayaraman D, Rougé P, Barre A, Lougnon G, Schornack S, Bono J-J, Cook DR. 2007. A novel nuclear protein interacts with the symbiotic DMI3 calcium- and calmodulin-dependent protein kinase of *Medicago truncatula*. Molecular Plant-Microbe Interactions 20(8): 912–921.

Murray JD, Muni RRD, Torres-Jerez I, Tang Y, Allen S, Andriankaja M, Li G, Laxmi A, Cheng X, Wen J. 2011. *Vapyrin,* a gene essential for intracellular progression of arbuscular mycorrhizal symbiosis, is also essential for infection by rhizobia in the nodule symbiosis of *Medicago truncatula*. The Plant Journal 65(2): 244–252.

Nadal M, Sawers R, Naseem S, Bassin B, Kulicke C, Sharman A, An G, An K, Ahern KR, Romag A. 2017. An N-acetylglucosamine transporter required for arbuscular mycorrhizal symbioses in rice and maize. Nature plants 3(6): 17073.

Oldroyd GED. 2013. Speak, friend, and enter: signalling systems that promote beneficial symbiotic associations in plants. Nature Reviews Microbiology 11: 252.

O’Connell RJ, Panstruga R. 2006. Tête à tête inside a plant cell: establishing compatibility between plants and biotrophic fungi and oomycetes. New Phytologist 171(4): 699–718.

Panstruga R. 2003. Establishing compatibility between plants and obligate biotrophic pathogens. Current opinion in plant biology 6(4): 320–326.

Parniske M. 2008. Arbuscular mycorrhiza: the mother of plant root endosymbioses. Nature Reviews Microbiology 6: 763.

Pavan S, Schiavulli A, Appiano M, Marcotrigiano AR, Cillo F, Visser RG, Bai Y, Lotti C, Ricciardi L. 2011. Pea powdery mildew *er1* resistance is associated to loss-of-function mutations at a *MLO* homologous locus. Theoretical and Applied Genetics 123(8): 1425–1431.

Pennington HG, Jones R, Kwon S, Bonciani G, Thieron H, Chandler T, Luong P, Morgan SN, Przydacz M, Bozkurt T. 2019. The fungal ribonuclease-like effector protein CSEP0064/BEC1054 represses plant immunity and interferes with degradation of host ribosomal RNA. PLoS pathogens 15(3): e1007620.

Piffanelli P, Zhou F, Casais C, Orme J, Jarosch B, Schaffrath U, Collins NC, Panstruga R, Schulze-Lefert P. 2002. The barley MLO modulator of defense and cell death is responsive to biotic and abiotic stress stimuli. Plant Physiology 129(3): 1076–1085.

Primrose SB, Twyman R. 2013. Principles of gene manipulation and genomics. Oxford, UK: John Wiley & Sons.

Pumplin N, Mondo SJ, Topp S, Starker CG, Gantt JS, Harrison MJ. 2010. *Medicago truncatula* Vapyrin is a novel protein required for arbuscular mycorrhizal symbiosis. The Plant Journal 61(3): 482–494.

Remy W, Taylor TN, Hass H, Kerp H. 1994. Four hundred-million-year-old vesicular arbuscular mycorrhizae. Proceedings of the National Academy of Sciences 91(25): 11841–11843.

Requena N, Voss S, Betz R, Heidt S, Corradi N. 2018. RiCRN1, a crinkler effector from the arbuscular mycorrhizal fungus Rhizophagus irregularis, functions in arbuscule development. Frontiers in microbiology 9: 2068.

Ruiz-Lozano JM, Gianinazzi S, Gianinazzi-Pearson V. 1999. Genes involved in resistance to powdery mildew in barley differentially modulate root colonization by the mycorrhizal fungus *Glomus mosseae*. Mycorrhiza 9(4): 237–240.

Saur IML, Bauer S, Lu X, Schulze-Lefert P. 2019. A cell death assay in barley and wheat protoplasts for identification and validation of matching pathogen AVR effector and plant NLR immune receptors. Plant methods 15(1): 118.

Tadege M, Wen J, He J, Tu H, Kwak Y, Eschstruth A, Cayrel A, Endre G, Zhao PX, Chabaud M, et al. 2008. Large-scale insertional mutagenesis using the *Tnt1* retrotransposon in the model legume *Medicago truncatula*. The Plant Journal 54(2): 335–347.

Takamatsu S. 2004. Phylogeny and evolution of the powdery mildew fungi (Erysiphales, Ascomycota) inferred from nuclear ribosomal DNA sequences. Mycoscience 45(2): 147–157.

Wang E, Schornack S, Marsh JF, Gobbato E, Schwessinger B, Eastmond P, Schultze M, Kamoun S, Oldroyd GE. 2012. A common signaling process that promotes mycorrhizal and oomycete colonization of plants. Current Biology 22(23): 2242–2246.

Wang Y, Cheng X, Shan Q, Zhang Y, Liu J, Gao C, Qiu J-L. 2014. Simultaneous editing of three homoeoalleles in hexaploid bread wheat confers heritable resistance to powdery mildew. Nature Biotechnology 32(9): 947.

Watts-Williams SJ, Cavagnaro TR. 2012. Arbuscular mycorrhizas modify tomato responses to soil zinc and phosphorus addition. Biology and Fertility of Soils 48(3): 285–294.

Xue L, Cui H, Buer B, Vijayakumar V, Delaux P-M, Junkermann S, Bucher M. 2015. Network of GRAS transcription factors involved in the control of arbuscule development in *Lotus japonicus*. Plant Physiology 167(3): 854–871.

Zhang Q, Blaylock LA, Harrison MJ. 2010. Two *Medicago truncatula* half-ABC transporters are essential for arbuscule development in arbuscular mycorrhizal symbiosis. The Plant Cell 22(5): 1483–1497.

Zhang X, Pumplin N, Ivanov S, Harrison MJ. 2015. EXO70I is required for development of a sub-domain of the periarbuscular membrane during arbuscular mycorrhizal symbiosis. Current Biology 25(16): 2189–2195.

