## Supporting information: Methods S1-S6, Table S1-S3, Figs. S1-S4 for "Mildew Locus O facilitates colonization by arbuscular mycorrhiza in angiosperms"

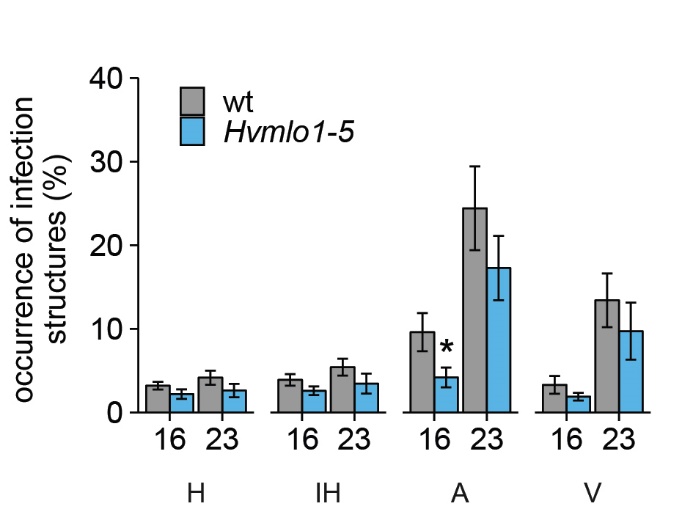

**Fig. S1 Mycorrhization in barley cv. Pallas wild type (wt) and *Hvmlo1-5* mutants.** Quantification of arbuscular mycorrhizal structures, hyphopodia (H), intraradical hyphae (IH), arbuscules (A), and vesicles (V) in wild type and *Hvmlo1-5* roots at 16 and 23 dpi with *Rhizophagus irregularis*. The binomial occurrences of mycorrhizal structures are shown as the percentage of the total number of root sections assessed. Statistical comparisons have been made to the wild type. Values are the mean of 12 biological replicates ±SEM (error bars) (General Linear Model with a logit link function; ANOVA; *, P < 0.05).

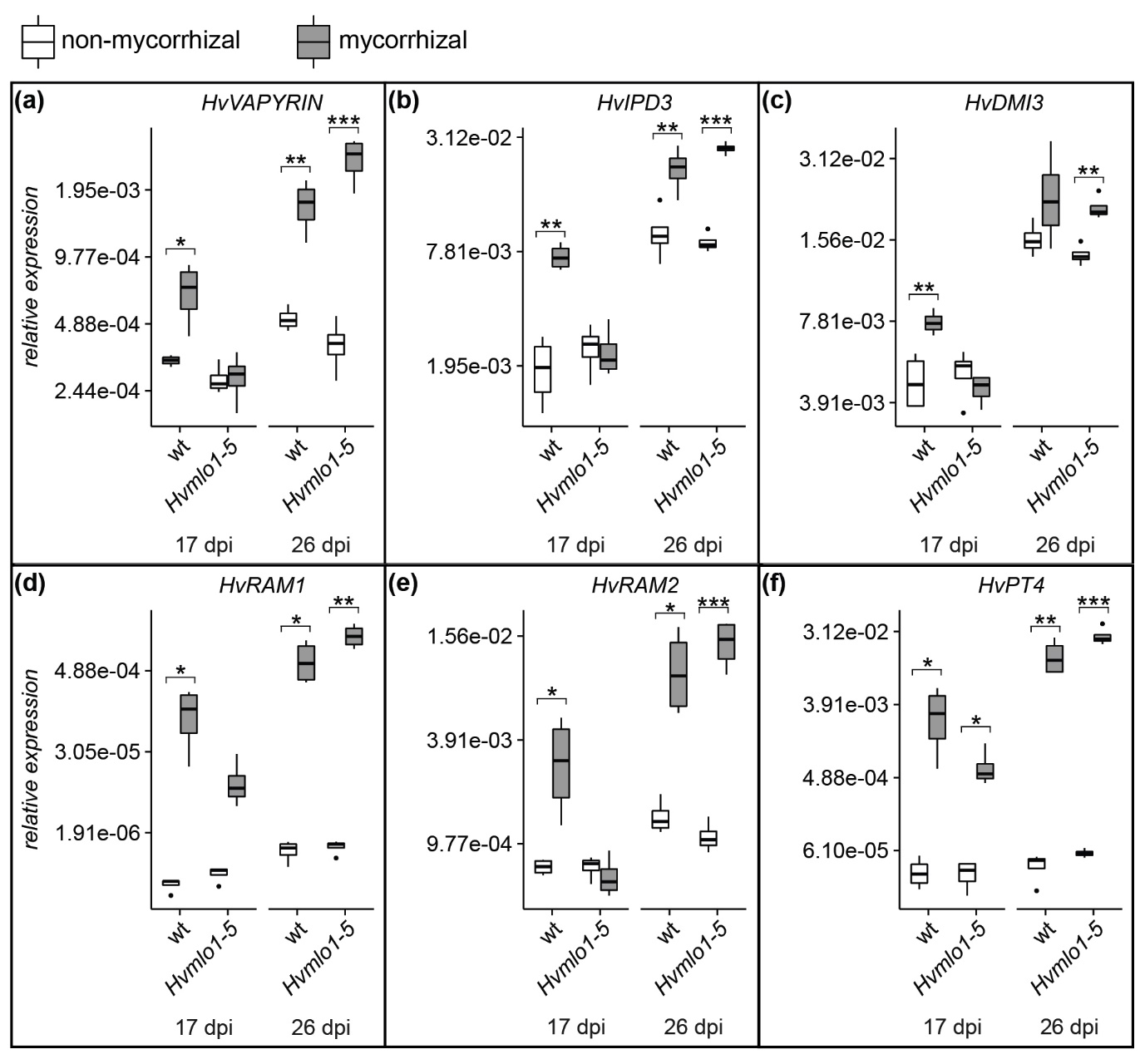

**Fig. S2 Relative expression of potential barley orthologues known mycorrhizal genes.** Relative expression of *VAPYRIN* (a), *IPD3* (b), *DMI3* (c), *RAM1* (d), *RAM*2 (e) and *PT4* (f) in barley cv. Ingrid wild type and *Hvmlo1-5* roots at 17 and 26 dpi with R. irregularis. Expression levels were measured by RT-qPCR and normalized to *HvEF1alpha*. Statistical comparisons were made relative to non-mycorrhized root samples. Bars represent means of 4 biological replicates ± SEM (error bars) (Student’s t-test: *, P < 0.05; **, P < 0.01; ***, P < 0.001).

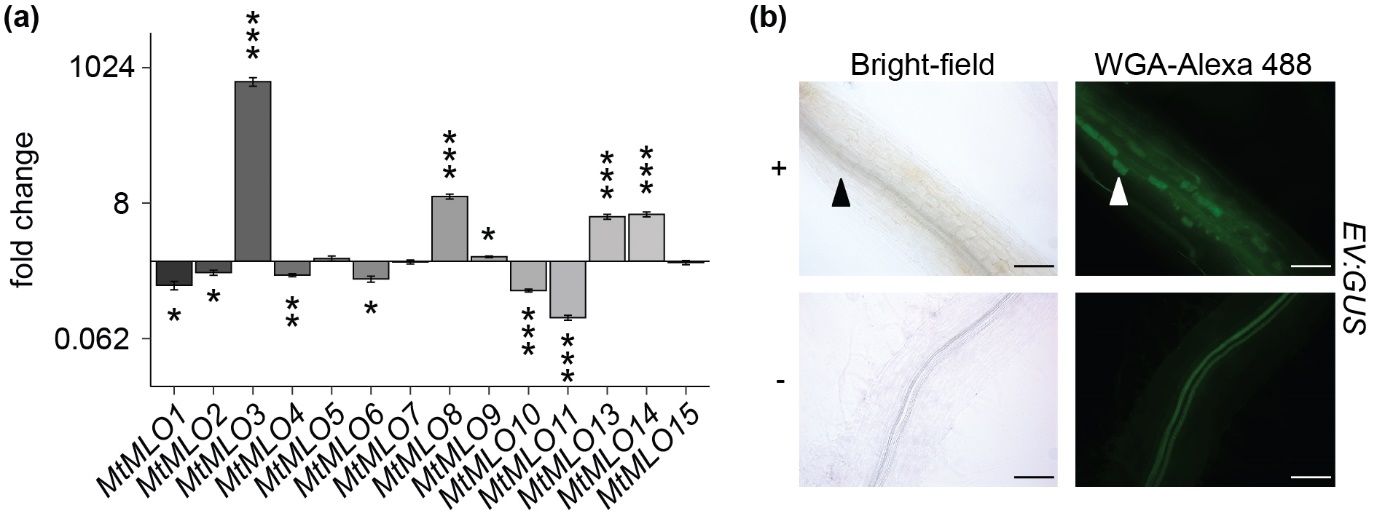

**Fig. S3 *MtMLO* gene expression.** (a) Quantification of induction of *MtMLO* genes during mycorrhization. Gene expression was measured by RT-qPCR in non-mycorrhizal versus mycorrhizal wild type *M. truncatula* cv. R108 roots at 22 dpi. Expression levels were measured by RT-qPCR and normalized to the geometric mean of *MtUBIQUITIN* and *MtPTB.* Statistical comparisons were made relative to non-mycorrhized root samples. Bars represent means of 8 biological replicates ± SEM (error bars) (Student’s t-test: *, P < 0.05; **, P < 0.01; ***, P < 0.001) (b) Activity of the empty vector control in mycorrhized and non‐mycorrhized wild type roots at 21 dpi. Bright-field and corresponding green fluorescence images of *M. truncatula* hairy roots transformed with an empty vector construct (no promoter driving the *β-glucuronidase (GUS)* gene). Mycorrhizal fungal structures were visualized using Alexa Fluor 488 wheat germ agglutinin. Solid arrowheads indicate cells containing arbuscules, and empty arrowheads indicate hyphopodia. Scale bars = 100 μm.

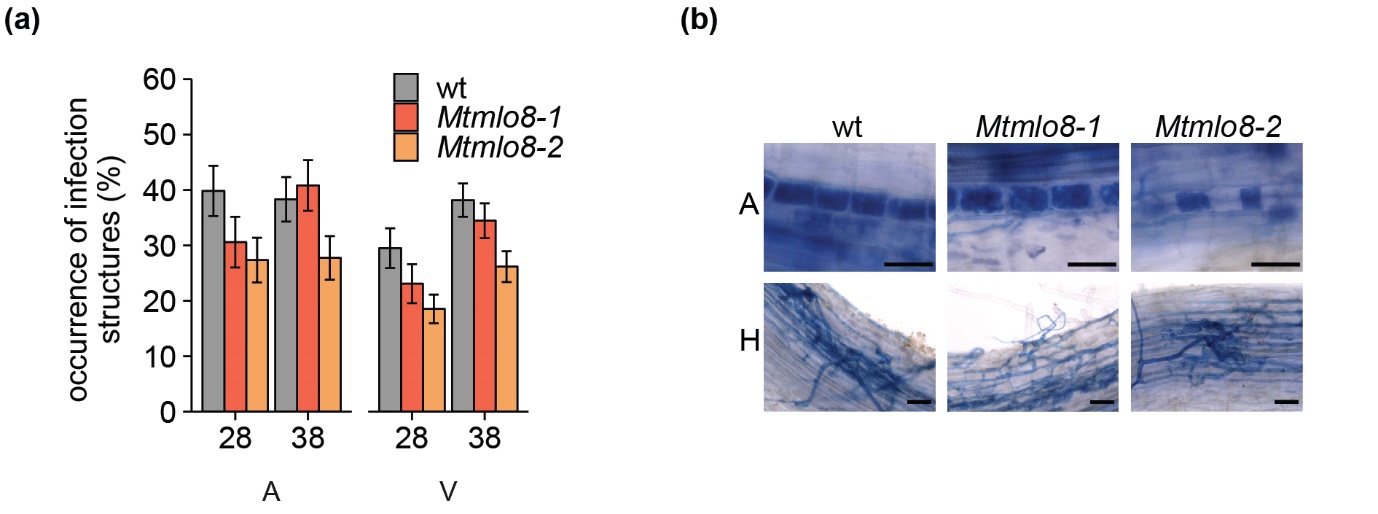

**Fig. S4 Mycorrhization in *M. truncatula* wild type and *Mtmlo8* mutants.** (a) Quantification of arbuscular mycorrhizal structures, arbuscules (A), and vesicles (V) in wild type and *Mtmlo8* roots at 28 and 38 dpi. The binomial occurrences of mycorrhizal structures are shown as a percentage of the total number of root sections assessed. Statistical comparisons have been made to the wild type. Values are the mean of 12 biological replicates ±SEM (error bars) (General Linear Model with a logit link function; ANOVA. (b) Appearance of hyphopodia (h) and arbuscules (A) in wild type and *Mtmlo8* mutant roots at 16 dpi with *R. irregularis*. Mycorrhizal fungal structures were visualized with ink-staining.

**Table S1** Normalized read counts for RNA-seq samples obtained from barley cv. Ingrid wild type and *Hvmlo1-5* roots at 17 dpi and 26 dpi with and without *R. irregularis*

Separate excel file – Table S1

**Table S2** Fold changes of RNA-seq samples obtained from barley cv. Ingrid wild type and *Hvmlo1-5* roots at 17 dpi and 26 dpi with and without R. *irregularis.* At each time point (17 dpi and 26 dpi), fold changes for all genes are shown for wild type: non-mycorrhizal versus mycorrhizal, *Hvmlo1-5:* non-mycorrhizal versus mycorrhizal, and mycorrhizal: wild type versus *Hvmlo1-5.*

Separate excel file – Table S2

**Table S3** Fold changes of potential orthologues of previously described mycorrhizal genes in non-mycorrhized versus mycorrhized wild type and *Hvmlo1-5* roots: ‘n.s.’ indicates FDR-corrected P-values > 0.05.

| ***Mt* gene** | ***Mt* identifier** | ***Hv* identifier** | **Fold change during mycorrhization** | | | |
| --- | --- | --- | --- | --- | --- | --- |
|  |  |  | **17 dpi** | | **26 dpi** | |
|  |  |  | **wt** | ***Hvmlo1-5*** | **wt** | ***Hvmlo1-5*** |
| *VAPYRIN* | Medtr6g027840 | HORVU4Hr1G002180 | 2.2 | 1.5 | 3.7 | 3.4 |
| *NSP1* | Medtr8g020840 | HORVU2Hr1G104160 | 4.6 | n.s. | n.s. | n.s. |
| *NSP2* | Medtr3g072710 | HORVU4Hr1G061310 | 4.4 | n.s. | 10.6 | 10.0 |
| *DMI1* | Medtr2g005870 | HORVU5Hr1G120340 | 1.7 | n.s. | 1.3 | n.s. |
| *DMI2* | Medtr5g030920 | HORVU2Hr1G058820 | 6.8 | 1.7 | 3.2 | 2.8 |
| *DMI3* | Medtr8g043970 | HORVU1Hr1G068660 | 1.9 | n.s. | n.s. | n.s. |
| *IPD3* | Medtr5g026850 | HORVU7Hr1G008420 | 4.9 | n.s. | 3.0 | 2.5 |
| *NOPE1* | Medtr3g093270  Medtr3g093290 | HORVU2Hr1G005470 | 6.6 | 1.9 | 6.6 | 9.5 |
| *RAD1* | Medtr4g104020 | HORVU3Hr1G088780 | 17.5 | n.s. | 11.5 | 9.8 |
| *RAM2* | Medtr1g040500 | HORVU4Hr1G011110 | 5.1 | n.s. | 10.2 | 8.1 |
| *EXO70I* | Medtr1g017910 | HORVU7Hr1G052100 | 144.4 | 31.7 | 467.5 | 303.0 |
| *STR* | Medtr8g107450 | HORVU5Hr1G060690 | 11.7 | 3.4 | 83.6 | 117.0 |
| *STR2* | Medtr5g030910 | HORVU3Hr1G066220 | 1048.7 | 61.6 | 1149.0 | 389.2 |
| *AMT2-3* | Medtr8g074750 | HORVU3Hr1G082610 | 3959.1 | 64.5 | 1719.0 | 1554.0 |
| *AMT2-4* | Medtr7g115050 | HORVU5Hr1G095030 | 312.2 | 153.5 | 2904.3 | 721.5 |
| *PT4* | Medtr8g074750 | HORVU6Hr1G058690 | 1238.1 | 268.1 | 4923.8 | 6312.1 |

**Methods S1** MLO protein sequences *from A. thaliana, A. trichopoda, B. vulgaris, D. caryophyllus, G. max, H. vulgare, L. angustifolius M. truncatula, O. sativa, P. patens, P. sativum, S. lycopersicum,* and *T. aestivum.*

Separate excel file – Methods S1

**Methods S2** **Gene structure of barley and wheat *MLO1*.** Arrows indicate (a) the X-ray-induced *(Hvmlo1-1)* and Ethyl methanesulfonate-induced *(Hvmlo1-5)* mutation sites in barley cv. Ingrid and Pallas *Hvmlo1* mutants (Jørgensen, 1992) and (b) the TALEN-induced mutation sites in the wheat cv. KN199 *Tamlo1* mutant (Wang et al., 2014).

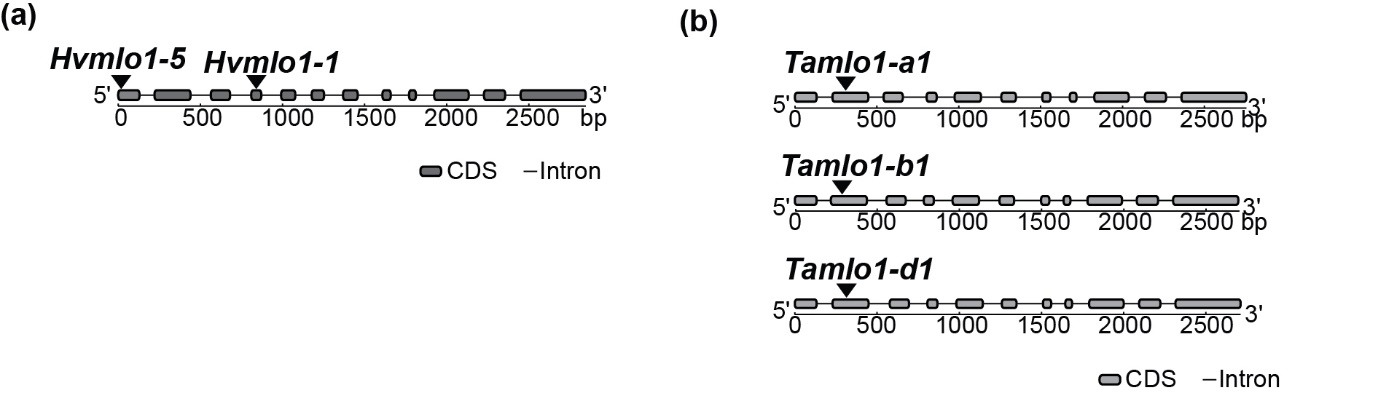

**Methods S3 Primer sequences**

| Description | Name, reference | Sequence |
| --- | --- | --- |
| **Barley** | | |
| RT-qPCR | HvEF1a_F (Schoonbeek *et al.*, 2015) | ATGATTCCCACCAAGCCCAT |
|  | HvEF1a_R (Schoonbeek *et al.*, 2015) | ACACCAACAGCCACAGTTTGC |
|  | HvMLO1_F | GCAAGCCCAGCAAGTACG |
|  | HvMLO1_R | GCGAGCACGAAGATGAAGAC |
|  | HvVAPYRIN_F | GTGGTGGATGTCCTCCTCAAG |
|  | HvVAPYRIN_R | CACCGACCTTCTCCAGTAACC |
|  | HvDMI3_F | GCTAGCAGAATTCGAGCAGGT |
|  | HvDMI3_R | CAGAGGATCTCCCTCATGTCC |
|  | HvIPD3_F | CGCAGAGCTTCAGAGAAGACA |
|  | HvIPD3_R | AGATGGTTGTCGAGGTCGATT |
|  | HvRAM1_F | CCAGGCTCCCAAGATTATGAC |
|  | HvRAM1_R | GAGTCGAAGATCGCCGAGTAG |
|  | HvRAM2_F | GGCTGGACCCCTTCTACTTCT |
|  | HvRAM2_R | AGGCGATGAGTCTCTGGATGT |
|  | HvPT4_F (Zhang *et al.*, 2010) | GGATTCTTTTGCACGTTCTTGG |
|  | HvPT4_R (Zhang *et al.*, 2010) | CCTGTCATTTGGTGTTGCAGTG |
| ***M. truncatula*** | | |
| Genotyping *Mtmlo8* mutants | TNT1 | CAGTGAACGAGCAGAACCTGTG |
|  | NF11523_F | TCACCTTTGCACTTGCTTCA |
|  | NF11523_R | AGAGCATGCCTTGGATGTGT |
|  | NF15162_F | CCATGAGTGAAGCCTTGGAG |
|  | NF15162_R | AAGGCACTTGACCCTGCATA |
| *MtMLO8* cDNA amplification | Full_MtMLO8_F | AGCTCCTGGGGAAAGAACAT |
|  | Full_MtMLO8_R | AAGGCTTATCAAATGAGAAGTCAATA |
| Gateway compatible primers (promoter-GUS) | attB_pMtMLO8_F | ggggacaagtttgtacaaaaaagcaggctTCTGGACCGGTTCAAAAATAAAA |
|  | attB_pMtMLO8_R | GAAAGAAGAAAGCAAATTAAGAacccagctttcttgtacaaagtggtcccc |
| RT-qPCR | MtUBIQUITIN_F (Kakar *et al.*, 2008) | GCAGATAGACACGCTGGGA |
|  | MtUBIQUITIN_R (Kakar *et al.*, 2008) | AACTCTTGGGCAGGCAATAA |
|  | MtPTB_F (Kakar *et al.*, 2008) | CGCCTTGTCAGCATTGATGTC |
|  | MtPTB_R (Kakar *et al.*, 2008) | TGAACCAGTGCCTGGAATCCT |
|  | MtMLO1_F  MtMLO1_R  MtMLO2_F  MtMLO2_R  MtMLO3_F  MtMLO3_R  MtMLO4_F  MtMLO4_R  MtMLO5_F  MtMLO5_R  MtMLO6_F  MtMLO6_R  MtMLO7_F  MtMLO7_R  MtMLO8_F  MtMLO8_R  MtMLO9_F  MtMLO9_R  MtMLO10_F  MtMLO10_R  MtMLO11_F  MtMLO11_R  MtMLO13_F  MtMLO13_R  MtMLO14_F  MtMLO14_R  MtMLO15_F  MtMLO15_R | TGCTTCCACAGAACAACTGC  CTGTGTGGTGCCATTTCTTG  AGACCCCTACTTGGGCTGTT  AATAATCCAATGGCCCATCA  AATTTGCATGGGAGTCTTCGT  TATGTTCTTCCTTGCGGTGTG  CTGAACGAGTGAAGCCATCAG  TCCATATGTGGACCAAATCCA  TGCAAAACTGGCGTAAAAATG  AGCTGTGTTATGCTCCTGCAA  AGCTTTTGGGATTGTGATGCT  GCCTTGGATGTTTGTTCATCA  TTGGTTTAGCGATGAGCAAGA  GGATGAGATGCATGATGGAAA  GACTATGCGGCATGGGTTTAT  GCAAAAGCCCATAAAGGAACA  TTCTGGATTGCTTTCGTTCCT  CTATGGCTGCATGCTTTTCAG  AGAAGATGGTGCCCCTTACAA  TGCCTTCCATCCTCGTATCTT  ATTCCTTTCCCGCACTTATGA  CAGTTTAGTCCCCGCAAGAAG  TTTTGGGTTGCTTTCATTCCT  TTTGCCTTGTATGGCTGAATG  TGTCAATGGTTGGCACACATA  TCACCTTGTATGGCTGAATGC  TTACACTTCGCAAAGGCTTCA  CCGACAACAAATCCCCATAGT |

**Methods S4 Sterilization methods and growth conditions**

| Figures | Species | Sterilization | Growth conditions |
| --- | --- | --- | --- |
| 1a,b | Barley | 2% sodium hypochlorite for 4 min, 5 x rinse sterile dH_2_O. Seeds were spread onto plates containing sterilized filter paper, then germinated for 2 days in darkness at 23°C | Germinated seedlings were transferred to pots containing 80% sterilized Terragreen/Sand (1:1 mix of terragreen (Oil‐dry UK ltd) and sharp sand (BB Minerals)) and 20% mycorrhizal inoculum (soil substrate containing *Allium schoenoprasum* roots exposed to *Rhizophagus irregularis* for 8 weeks). Plants were grown in a glasshouse with no additional heat or light during June – August 2018 (temperature range: 14°C - 48°C; average temperature: 28°C). |
| 1c,d,e | Barley |  | Germinated seedlings were transferred plates containing modified Fahraeus plant agar medium (ModFP) with Augmentin (50 μg/ml), grown in a controlled-environment room at 23°C (16-h photoperiod, and 300 mmol m^-2^ s^-1^) for 3 days, then transferred to 90% sterilized Terragreen/Sand and 10% mycorrhizal inoculum (soil substrate containing *Allium schoenoprasum* roots exposed to *Rhizophagus irregularis* for 8 weeks). Plants were grown in a Weiss Technik growth cabinet at 35°C/20°C (35% RH day, 50 % RH night; 16-h photoperiod, and 500 μmol m^-2^ s^-1^, average temperature: 30°C) and harvested at 17 dpi and 26 dpi. |
| 1f, S1 | wheat, barley |  | Germinated seedlings were transferred to pots containing 90% sterilized Terragreen/Sand (1:1 mix of terragreen (Oil‐dry UK ltd) and sharp sand (BB Minerals)) and 10% mycorrhizal inoculum (soil substrate containing *A. schoenoprasum* roots exposed to *Rhizophagus irregularis* for 8 weeks). Plants were grown in a glasshouse with no additional heat or light during June – August 2018 (temperature range: 14°C - 48°C; average temperature: 28°C). |
| 3a,e, S4 | *M. truncatula* | Concentrated sulphuric acid for 8 min, 5 x rinse sterile dH_2_O, 10% sodium hypochlorite for 4 min, 5 x rinse sterile dH_2_O. Seeds were imbibed in sterile dH_2_O containing Nystatin (5 μg/ml) and Augmentin (50 μg/ml) for 5 h, then placed onto water agar containing Nystatin (5 μg/ml) and Augmentin (50 μg/ml). Seeds were stratified for 5 days in darkness at 4°C, then seeds germinated overnight in darkness at 23°C. | Germinated seedlings were grown on plates containing modFP in a controlled-environment room for 7 days at 23°C (16-h photoperiod, and 300 mmol m^-2^ s^-1^) then transferred to pots containing 90% sterilized Terragreen/Sand and 10% mycorrhizal inoculum. Plants were grown in a controlled environment room at 22°C (80% humidity, 16-h photoperiod, and 300 mmol m^-2^ s^-1^). |
| 3b,f | *M. truncatula* |  | Roots of germinated seedlings were transformed by *Agrobacterium rhizogenes*-mediated gene transfer (Boisson-Dernier *et al.*, 2001) and grown for 3 weeks on plates containing modFP in a controlled-environment room at 23°C (16-h photoperiod, and 300 mmol m^-2^ s^-1^). Seedlings were transferred to pots containing 90% sterilized Terragreen/Sand and 10% mycorrhizal inoculum. Plants were grown in a controlled environment room at 22°C (80% humidity, 16-h photoperiod, and 300 mmol m^-2^ s^-1^). |
| 3g | *M. truncatula* |  | Germinated seedlings were transferred to soil and grown for 3 weeks in a Schneider growth cabinet at 18°C (16-h photoperiod 300 mmol m^-2^ s^-1^). Detached *M. truncatula* leaves were placed on distilled water agar plates containing benzimidazole (10 μg/ml) and inoculated with by blowing fresh *Erysiphe pisi* (Ep) CJ001 spores into an inoculation tower. |

**Methods S5 Staining and visualization methods**

| Figures | Description | Staining | Visualization |
| --- | --- | --- | --- |
| 1b,d,f, 3e,f, S1, S4 | Ink-staining of mycorrhizal structures | Roots were washed in tap water and then incubated in 10% KOH for 5 min at 96 °C, rinsed in dH_2_O, then stained using 5% black ink (Waterman) and 5% acetic acid for 3 min at 96 °C. Roots were de-stained in dH_2_O for 1 day. | M80 microscope (Leica) |
| 1c, 3b | Fluorescence staining of mycorrhiza­l structures | Roots were washed in sterile dH_2_O, placed in 50% ethanol overnight at room temperature, then placed in 20% KOH for 2 days at room temperature. Root samples were rinsed in sterile dH2O, incubated in 0.1M HCL for 3 h, rinsed in sterile dH2O, then rinsed in 1 x phosphate-buffered saline (PBS). Root samples were stained overnight with Wheat Germ Agglutinin (WGA) labeled with Alexa Fluor 488 by incubation in a 1 x PBS solution containing 0.4 μg/ml Alexa Fluor 488 WGA in darkness at 23°C. Samples were de-stained and stored in 1 x PBS at 4°C. | DM 6000 microscope (Leica) and a DFC420 colour camera (Leica). |
| 3b | GUS-staining | Roots were washed in sterile dH_2_O then 1 x phosphate buffer pH 7.0 (NaH_2_PO_4_-Na_2_HPO_4_). Roots were vacuum infiltrated for 30 min in GUS staining solution (50 mM phosphate buffer pH 7.0, 0.5 mM K_3_Fe(CN)_6_, 0.5 mM K_4_Fe(CN)_6_, 50 mM EDTA, 3% sucrose and 2 mM X-Gluc (5-bromo-4-chloro-3-indolyl-beta-D-glucuronide), incubated in the dark at 37°C for 2 h, then rinsed in 20% ethanol for 20 mins, 50% ethanol for 20 mins, and 70% ethanol for 20 mins. | DM 6000 microscope (Leica) and a DFC420 colour camera (Leica). |
| 3d | Trypan blue staining of  powdery mildew | Leaves were placed in 70% ethanol and agitated at 150 rpm. After 6 h, leaves were placed in 100% ethanol and agitated at 150 rpm. Ethanol was regularly changed until chlorophyll was removed, and the leaves were white. Leaves were placed in lactoglycerol (1:1:1 lactic acid: glycerol: dH2O) for 30 min, then stained using Trypan blue stain (0.1% Trypan blue in lactoglycerol) for 10 min. | Vickers microscope |

**Methods S6 Gene expression analyses. Methods for RT-qPCR and transcriptome analyses**

For RT-qPCR, 150 ng of RNA was retrotranscribed (SuperScript IV reverse transcriptase, Invitrogen). For RT-qPCR analysis, gene expression was monitored by SYBR Green-based quantitative PCR using a LightCycler LC480 system using gene-specific primers (Methods S3). Data were analyzed according to the 2^-ΔΔCT^ method (Livak & Schmittgen, 2001) using amplification efficiency corrections.

For transcriptome analysis, RNA sequencing was performed by Novogene (Cambridge, UK). mRNA libraries were prepared with the Illumina TruSeq® Stranded mRNA HT technology with a paired-end 150 bp (PE 150) strategy. To obtain pseudo-counts, Kallisto (Bray et al., 2016) version 0.44 was used. Coding sequences (CDS) from the genome of barley cv. Morex (Beier et al., 2017) were used as reference. For subsequent analysis, expression values from all splice forms were combined to one expression value per gene. Differential gene expression analysis was performed using DEGUST v.3.2.0 (Powell et al., 2019) and edgeR was used as the normalization method. Genes with an assigned false discovery rate (FDR)-corrected P-value >0.05 were discarded from further analysis. Normalized read counts for all treatments can be found in Table S1 and are deposited on the Gene Expression Omnibus (GEO) database.
